# Ring-shaped multimeric structure enables the acceleration of KaiB-KaiC complex formation induced by the ADP/ATP exchange inhibition

**DOI:** 10.1101/2021.07.06.451233

**Authors:** Shin-ichi Koda, Shinji Saito

## Abstract

Circadian clocks tick a rhythm with a nearly 24-hour period in various organisms. The clock proteins of cyanobacteria, KaiA, KaiB, and KaiC, compose a minimum circadian clock. The slow KaiB-KaiC complex formation, which is essential in determining the clock period, occurs when the C1 domain of KaiC binds ADP produced by ATP hydrolysis. KaiC is considered to promote this complex formation by inhibiting the backward process, ADP/ATP exchange, rather than activating the forward process, ATP hydrolysis. Remarkably, although inhibition of backward process, in general, decelerates the whole process, KaiC oppositely accelerates the complex formation. In this article, by building a novel reaction model, we investigate the molecular mechanism of the simultaneous promotion and acceleration of the complex formation, which may play a significant role in keeping the period invariant under environmental perturbations. Based on several experimental results, we assume in this model that six KaiB monomers cooperatively and rapidly binds to C1 with the stabilization of the binding-competent conformation of C1 only when C1 binds six ADP. We find the cooperative KaiB binding effectively separates the pre-binding process of C1 into a fast transformation to binding-competent C1 requiring multiple ATP hydrolyses and its slow backward transformation. Since the ADP/ATP exchange retards the forward process, its inhibition results in the acceleration of the complex formation. We also find that, in a simplified monomeric model where KaiB binds to a KaiC monomer independently of the other monomers, the ADP/ATP exchange inhibition cannot accelerate the complex formation. In summary, we conclude that the ring-shaped hexameric form of KaiC enables the acceleration of the complex formation induced by the backward process inhibition because the cooperative KaiB binding arises from the structure of KaiC.

**Author summary:** Circadian clocks tick a rhythm with a nearly 24-hour period in various organisms. The cyanobacterial circadian clock has attracted much attention because of its simplicity, composed of only three proteins, KaiA, KaiB, and KaiC. The rate of the slow KaiB-KaiC complex formation, which plays an essential role in the period determination, is mainly regulated by the ATP hydrolysis (forward process) and the ADP/ATP exchange (backward process) of KaiC. KaiC promotes the complex formation by inhibiting the backward process rather than activating the forward process. Remarkably, although inhibition of backward process, in general, slows down the whole process, KaiC oppositely accelerates the complex formation. In this article, we investigate the molecular mechanism of this acceleration by building a novel mathematical model based on several significant experimental results. We find the cooperative binding of six KaiB to a KaiC hexamer, which arises from the ring-shaped hexameric structure of KaiC, effectively separates the pre-binding process into a fast transformation to the binding-competent KaiC requiring multiple ATP hydrolyses and its slow backward transformation. Since the ADP/ATP exchange retards the forward process, its inhibition results in the acceleration of the complex formation.

## Introduction

Circadian clocks are endogenous timing systems. Organisms ranging from bacteria to higher plants and animals use the clocks to adapt their activity to daily changes in the environment. The cyanobacterial circadian clock has attracted much attention because it oscillates without transcription-translation feedback [1] and can be reconstituted *in vitro* by mixing three proteins, KaiA, KaiB, and KaiC, in the presence of ATP [2].

KaiC, the main component of the oscillator, forms a homohexamer consisting of N-terminal C1 and C-terminal C2 rings [3, 4]. In the presence of KaiA and KaiB, two amino acid residues near the ATP binding site in C2, Ser431 and Thr432, are periodically phosphorylated and dephosphorylated [5, 6]. During this oscillation, KaiA facilitates the phosphorylation by acting on C2 [7–10]. After C2 is phosphorylated, KaiB, on the other hand, inhibits the KaiA activity on C2 [11, 12] by forming the stable C1-KaiB-KaiA complex [13–15]. Then, C2 returns to the unphosphorylated state.

Keeping period constant is a fundamental function of biological clocks. For understanding the period-determining mechanism, slow processes of the KaiABC oscillator have intensively been studied. For example, the binding of KaiB to C1, taking a few hours, affects the period of the KaiABC oscillator as well as that of the damped oscillation in the absence of KaiA *in vivo* [16]. This KaiB-C1 complex formation proceeds in a dual conformational selection manner, i.e., both KaiB and C1 need to change their conformations into binding-competent ones before the binding. Thus, there have been attempts to determine the slower process of the two. An experimental study has reported a slow KaiB-KaiC complex formation [17]. This study has explained the slowness by assuming a slow conformational transition of KaiB. In contrast to this experiment conducted at only one protein concentration, another experiment has measured the KaiB concentration dependence of the binding rate [18]. Remarkably, in light of the kinetic theory [19], the result indicates that the pre-binding conformational transition of KaiC is the rate limiting process. Moreover, we have theoretically shown that the slow KaiB-KaiC binding of the former experiment can be explained by the attractive KaiB-KaiB interaction in the ring-shaped KaiB-KaiC complex [15], without assuming the slow conformational transition of KaiB [20]. Thus, it is now conceivable that the slowness of the binding arises from KaiC rather than KaiB.

To understand the whole picture of the KaiB-KaiC complex formation, it is then necessary to reveal the detail of the pre-binding processes of KaiC. Several experiments have shown that the ATPase activity of C1 closely relates to the complex formation. For example, a mutation disabling the ATP hydrolysis of C1 [21] or introduction of a non-hydrolyzable ATP analogue inhibits the complex formation [18, 21]. Moreover, the binding-competent conformation of KaiC binds ADP in C1 [14, 15, 22]. These results imply that the conformational transition to the binding-competent conformation occurs after the ATP hydrolysis of C1. This picture of the KaiC’s pre-binding process can be briefly summarized as

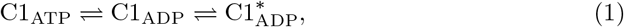

where the subscripts represent the nucleotide binding to C1, and the asterisk indicates the binding-competent conformation. This type of pre-binding process involving the ATPase activity can be seen in a molecularly detailed reaction model [23], for example.

In the scheme in Eq (1), the population of the binding-competent conformation is considered to be controlled by the exchange rate of bound ADP for exogenous ATP rather than the rate of ATP hydrolysis (the backward and forward processes of C1_ATP_ ⇌ C1_ADP_, respectively), i.e., the binding-competent C1 increases by inhibiting the ADP/ATP exchange [23]. This mechanism is consistent with the experimental result that a phosphomimetic mutant of KaiC with a high KaiB affinity, S431D/T432E-KaiC (KaiC-DE), has a low ATPase activity [24] because the ADP/ATP exchange inhibition reduces the ATPase activity. To sum up, it is conceivable that the phosphorylation of C2 shifts the equilibrium of Eq (1) toward the forward direction by inhibiting the backward process (that is, the ADP/ATP exchange of C1).

In the present article, we focus on the binding rate of KaiB to C1, which determines the time required for the complex formation. A recent experiment reported that S431E/T432E-KaiC (KaiC-EE), which may have a lowered ADP/ATP exchange rate of C1 as in KaiC-DE above, shows a faster KaiB-KaiC complex formation than the wild-type KaiC (KaiC-WT) [16]. That is, the more the complex formation is promoted due to the phosphorylation of C2, the more accelerated. This property may have a significant role in the period robustness against environmental perturbation such as temperature because it can prevent excess phosphorylation of C2 from taking time by accelerating the complex formation, which terminates the phosphorylation. Thus, the molecular mechanism of this simultaneous promotion and acceleration of the KaiB-KaiC complex formation is worth investigating in detail. However, the simple scheme in Eq (1) cannot explain it. Under this scheme, the ADP/ATP exchange inhibition, which promotes the binding, reduces the apparent binding rate because the relaxation rate of C1_ATP_ ⇌ C1_ADP_ is given by the sum of the rates of ATP hydrolysis and ADP/ATP exchange. This contradiction with the experimental result implies that the scheme in Eq (1) is oversimplified or lacks some important factors.

In the following section, we propose a reaction model of the KaiB-KaiC complex formation with pre-binding processes of KaiC, which is an extension of the scheme in Eq (1). The present model explicitly considers the hexameric form of KaiC and imposes several restrictive conditions for the binding. Specifically, reflecting that KaiB_6_KaiC_6_ complex is much more stable than KaiB_*n*_KaiC_6_ (*n <* 6) due to the strong attractive adjacent KaiB-KaiB interaction [15, 20, 25, 26], the present model assumes that six KaiB monomers immediately bind to C1 only when C1 binds six ADP. Moreover, the ADP/ATP exchange rate of the binding-competent C1 is assumed to be smaller than that of the binding-incompetent C1. Based on these assumptions, we show that the present hexameric model can explain the acceleration of the KaiB-KaiC complex formation by the ADP/ATP exchange inhibition, whereas a simplification to monomeric model fails to explain. This result suggests that the hexameric ring-shaped form of KaiC and the attractive adjacent KaiB-KaiB interaction are essential factors for accelerating the complex formation. As a corollary, we further show that the binding of KaiB to C1 reduces the ATPase activity of C1 [24], which has not been explained by any mathematical model so far.

## Results

### Model setup

In this study, we build a reaction model of the binding of KaiB to the C1 ring of a KaiC hexamer. The present model explicitly considers the nucleotides binding to C1. We denote KaiC that binds *n* ATP and (6 − *n*) ADP in C1 by C_6,*n*_(0 ≤ *n* ≤ 6). Because the ATP binding sites of KaiC are almost always occupied by ATP or ADP [27], we ignore KaiC that binds less than six nucleotides in C1. We consider the ATP hydrolysis as a transition from C_6,*n*_ to C_6,*n*−1_ and the ADP/ATP exchange as the backward process. We ignore the ATP synthesis due to its high activation energy.

In the present model, we impose the following five assumptions on the KaiB-KaiC complex formation.

- (i) The C1 domain of each monomer in a KaiC hexamer has binding-competent and binding-incompetent conformations, and the conformational transition between them undergoes independently of the other monomers.
- (ii) The conformational transition to the binding-competent conformation is feasible only in a monomer binding ADP.
- (iii) The binding-competent conformation is negligibly populated in the absence of KaiB but is stabilized by the binding of KaiB.
- (iv) The complexes between a KaiC hexamer and less than six KaiB monomers are negligible, while KaiB6KaiC6 complex is highly stable.
- (v) The conformational transitions of C1 and KaiB and the association/dissociation of KaiB to/from KaiC are much faster than the ATP hydrolysis and the ADP/ATP exchange of C1.

These five assumptions are supported by the following reasons. Assumption (i) is consistent with the experimental result that several crystal structures of the C1 ring consist of multiple conformations of the monomer [22], suggesting that each monomer is not strongly constrained by the other monomers. Although there can still be weak cooperativity in C1, we assume the complete independence in the present model for simplicity. Assumption (ii), which is also adopted by the scheme in Eq (1), is based on the experimental result that the KaiB-KaiC complex binds ADP in C1 [14, 15, 22]. Assumption (iii) is consistent with the experimental result that the addition of KaiB increases the binding-competent conformation of C1 [18]. Although a small amount of the binding-competent conformation may exist in the real system even in the absence of KaiB, we ignore it for simplicity. Note that this assumption is a typical case of the conformational-selection binding mechanism, where the binding stabilizes a less populated energetically excited conformation. Assumption (iv) is based on the experimental observations on the stoichiometry of the KaiB-KaiC complex [25, 26], where the adjacent KaiB-KaiB interaction stabilizes the complex [15, 20]. Lastly, Assumption (v) is consistent with the experimental result that the ATPase activity of C1 affects the period [22, 24], which implies that the other processes involved in the KaiB-KaiC binding are fast. Moreover, the conformational transition of KaiB is theoretically shown to be not necessary slow [20].

With these assumptions, we construct the present model as follows. In the absence of KaiB, due to Assumptions (ii) and (iii), KaiC takes only the binding-incompetent conformation in the model. Thus, in this case, the present model consists of the ATP hydrolysis and the ADP/ATP exchange of C1 among binding-incompetent KaiC, C_6,*n*_ (0 ≤ *n* ≤ 6). Explicitly, the model is represented as

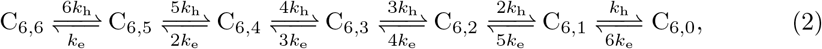

where *k*_h_ and *k*_e_ are the rate constants of the ATP hydrolysis and the ADP/ATP exchange of a binding-incompetent C1 monomer, respectively.

In the presence of KaiB, KaiB binds to C1 cooperatively, strongly, rapidly, and only when C1 binds six ADP

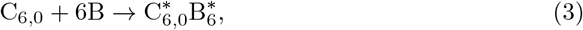

where B and B* represents a binding-incompetent and binding-competent KaiB monomers, respectively, and 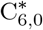 is a KaiC hexamer consisting of six binding-competent monomers. The strong cooperative binding of six KaiB, where its backward process can be disregarded, arises from Assumption (iv). This cooperative binding occurs only when C1 binds six ADP because all the six KaiC monomers in a KaiC hexamer need to bind ADP to take the binding-competent conformation due to Assumptions (i) and (ii). Assumption (v) requires the conformational transitions of KaiB and KaiC and the KaiB-KaiC binding to be fast. For simplicity, we assume that the process of Eq (3) immediately proceeds once C_6,0_ is produced in the presence of abundant KaiB. On the other hand, we also assume that 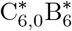 immediately dissociates once a bound ADP is exchanged into ATP because KaiC with less than six ADP in C1 cannot form a complex in the present model.

To sum up, the whole binding scheme in the presence of abundant KaiB is given by

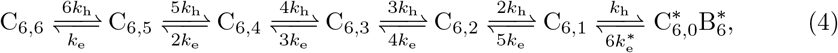

where 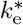 is the rate constant of the ADP/ATP exchange of a binding-competent C1 monomer. Eq (4) differs from Eq (2) in the last transition between C_6,1_ and 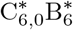, which represents the immediate formation and dissociation of the KaiB-KaiC complex mentioned above. The rates of them are dominated by the ATP hydrolysis and the ADP/ATP exchange of C1, respectively.

Moreover, to clarify the roles of the hexameric form of KaiC, we consider simplified monomeric schemes:

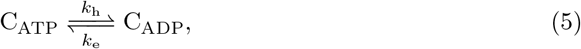

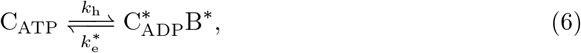

where C_ATP_ and C_ADP_ are KaiC monomers that bind ATP and ADP in C1, respectively. Note that this monomeric scheme can be derived if Assumption (iv) is modified as follows.

- (iv)’ A KaiB monomer can bind to a binding-competent C1 monomer independent of the other monomers.

Thus, the only difference between the hexameric and the monomeric schemes is the cooperativity of the KaiB binding originating from the attractive adjacent KaiB-KaiB interaction.

### Overview of the KaiB-KaiC complex formation

We here add Assumption (vi) on *k*_e_ and 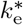;

- (vi) 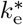 is smaller than *k*_e_.

With this assumption and the schemes in Eqs (2) and (4) (or Eqs (5) and (6) for the monomeric model), the KaiB-KaiC complex formation proceeds along with the following scenario. When abundant KaiB is added to a KaiC only system equilibrated according to Eq (2), KaiB immediately binds to initially existing C_6,0_. Then, 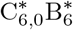 gradually increases because the relation 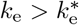 shifts the equilibrium toward 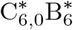 in Eq (4). This initially rapid and subsequently slow binding is consistent with the experimental observation [16]. Note that the apparent binding rate is given by the relaxation rate of the scheme in Eq (4).

### Acceleration of the binding by inhibiting the ADP/ATP exchange

As mentioned in Introduction, the KaiB-KaiC complex formation is considered to be controlled by changing the ADP/ATP exchange rate of C1 rather than the ATP hydrolysis. Specifically, the inhibition of the ADP/ATP exchange of C1 leads to the accumulation of ADP in C1 and then increases the binding-competent C1. In the present model, lowering *k*_e_ and/or 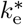 shifts the equilibrium of Eq (4) toward 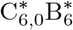. On the other hand, we focus on the experimental result that the apparent binding rate of KaiB to KaiC-EE is larger than KaiC-WT, suggesting that the ADP/ATP exchange inhibition accelerates the complex formation [16]. In the following, we show that while the monomeric binding scheme in Eq (6) cannot simultaneously achieve the promotion and acceleration of the complex formation by lowering the ADP/ATP exchange rate, the multimeric binding scheme in Eq (4) can.

To simulate the behaviors of KaiC-WT and KaiC-EE by the present model, we consider a set of rate constants, *k*_h_, *k*_eWT_, 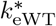, *k*_eEE_, and 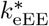, where *k*_h_ is the ATP hydrolysis rate constant common to KaiC-WT and KaiC-EE, and *k*_eWT(EE)_, 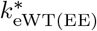 are the ADP/ATP exchange rate constants of the binding-incompetent and binding-competent KaiC-WT (KaiC-EE), respectively. Because C2 may affect the ADP/ATP exchange rate of C1 rather than the ATP hydrolysis rate, we assume that *k*_h_ is common to KaiC-WT and KaiC-EE for simplicity.

First, we examine whether the present model can reproduce the experimental result of the KaiB-KaiC complex formation (Fig 1). We calculate the amount of KaiB binding to KaiC-WT with (*k*_h_, *k*_eWT_, 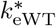) and to KaiC-EE with (*k*_h_, *k*_eEE_, 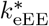) along with the hexameric scheme in Eq (4) and the monomeric scheme in Eq (6). The initial states are set to be the steady states of Eqs (2) and (5), respectively.

**Fig 1.**
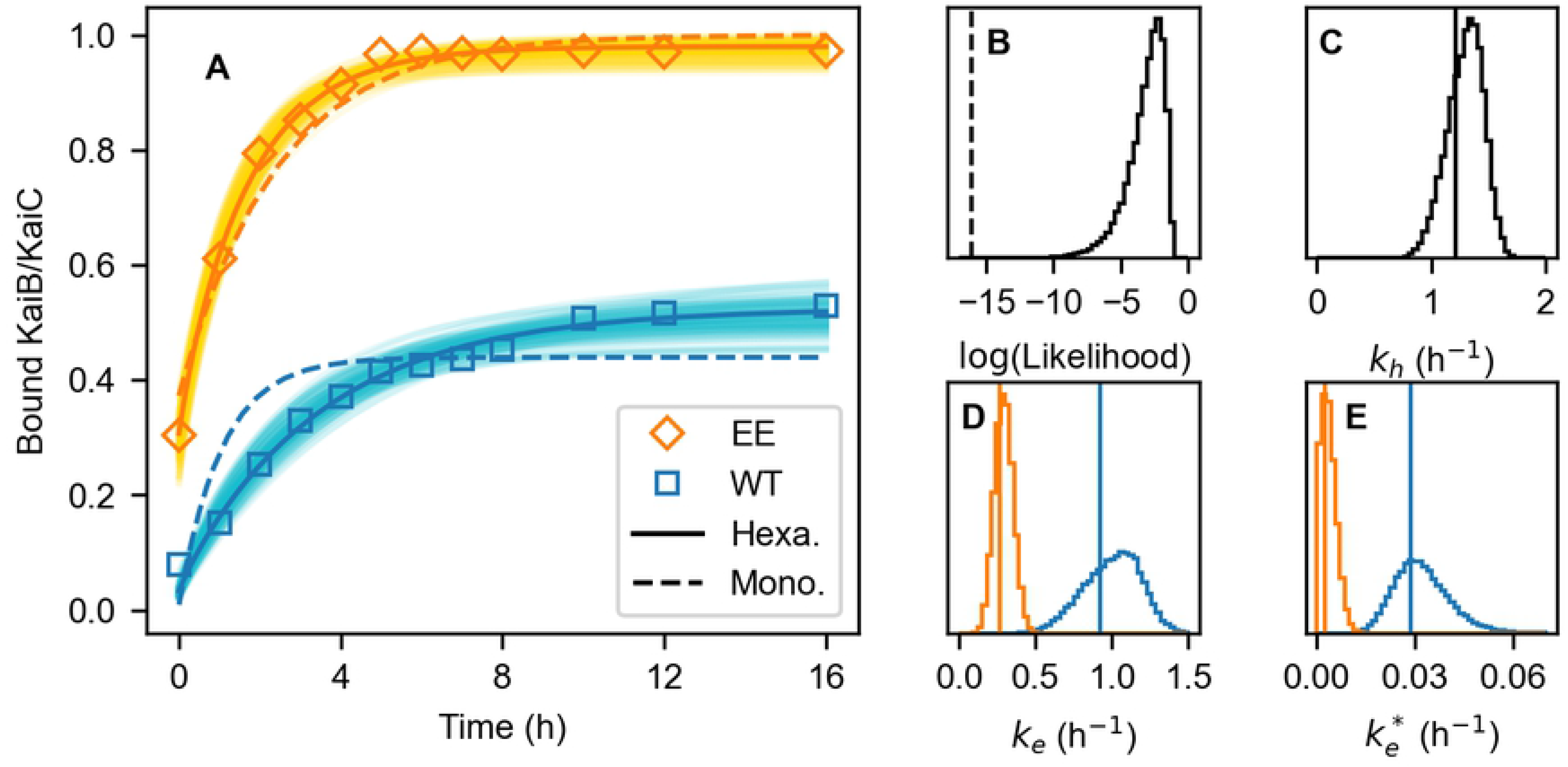
Results of the Bayesian parameter estimation. (A) Time courses of the binding of KaiB to KaiC-WT (blue) and KaiC-EE (orange). Squares are the experimental data. Bold solid and dashed curves are the best fits of the hexameric and monomeric models, respectively. Thin solid curves show 100 results of the hexameric model randomly chosen from the MCMC sampling. (B) Distribution of the hexameric model’s likelihood logarithm, which is defined in Methods. The vertical dashed line shows the likelihood logarithm of the monomeric model’s best fit. (C-E) Distributions of the hexameric model’s rate constants: (C) ATP hydrolysis *k*_h_, (D) ADP/ATP exchange of the binding-incompetent conformation *k*_e_ of KaiC-WT (blue) and KaiC-EE (orange), and (E) ADP/ATP exchange of the binding-competent conformation 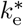 of KaiC-WT (blue) and KaiC-EE (orange). Vertical lines show the corresponding best-fit values.

Since the hexameric model has a wide range of the rate constants showing good fits (Fig 1A), we sample their distributions by Bayesian parameter estimation with Markov chain Monte Carlo (Figs 1C-E). In this sampling, to narrow the range of the parameters, we additionally use the experimental data of the ATPase activities of KaiC in the presence and absence of KaiB (see next subsection and Methods). This result shows that both *k*_eEE_ and 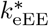 are lowered compared to *k*_eWT_ and 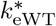, respectively (Figs 1D and E), indicating that the hexameric model can simultaneously achieve the promotion and acceleration of the KaiB-KaiC complex formation by lowering *k*_e_ and 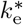.

On the other hand, the monomeric model fails to reproduce the KaiB bindings to KaiC-WT and KaiC-EE simultaneously (Figs 1A and B). This best fit only reproduces the promotion of the binding by inhibiting the ADP/ATP exchange and cannot reproduce the acceleration of the binding. Thus, the monomeric model cannot accelerate the binding by inhibiting the ADP/ATP exchange.

Next, we analyze the binding rate in detail. As mentioned in the previous subsection, because of the rapid binding, the apparent binding rate is given by the relaxation rate, that is, the smallest non-zero eigenvalue of the transition matrix of the pre-binding process shown in Eq (4) or (6). We denote them by *γ*_hexa_ and *γ*_mono_, respectively.

In the monomeric scheme, *γ*_mono_ is given by the sum of the rate constants of the forward and backward processes as

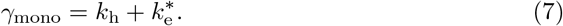

Thus, *γ*_mono_ never increases by lowering *k*_e_ or 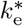. This is why the monomeric model cannot accelerate the binding by inhibiting the backward process.

In the hexameric scheme, any analytical expression of *γ*_hexa_ is not available. Instead, we numerically calculate *γ*_hexa_ as a function of *k*_e_ and 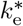 (Figs 2A and B). This result indicates that *γ*_hexa_ is an increasing function with respect to 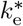 as well as *γ*_mono_.

**Fig 2.**
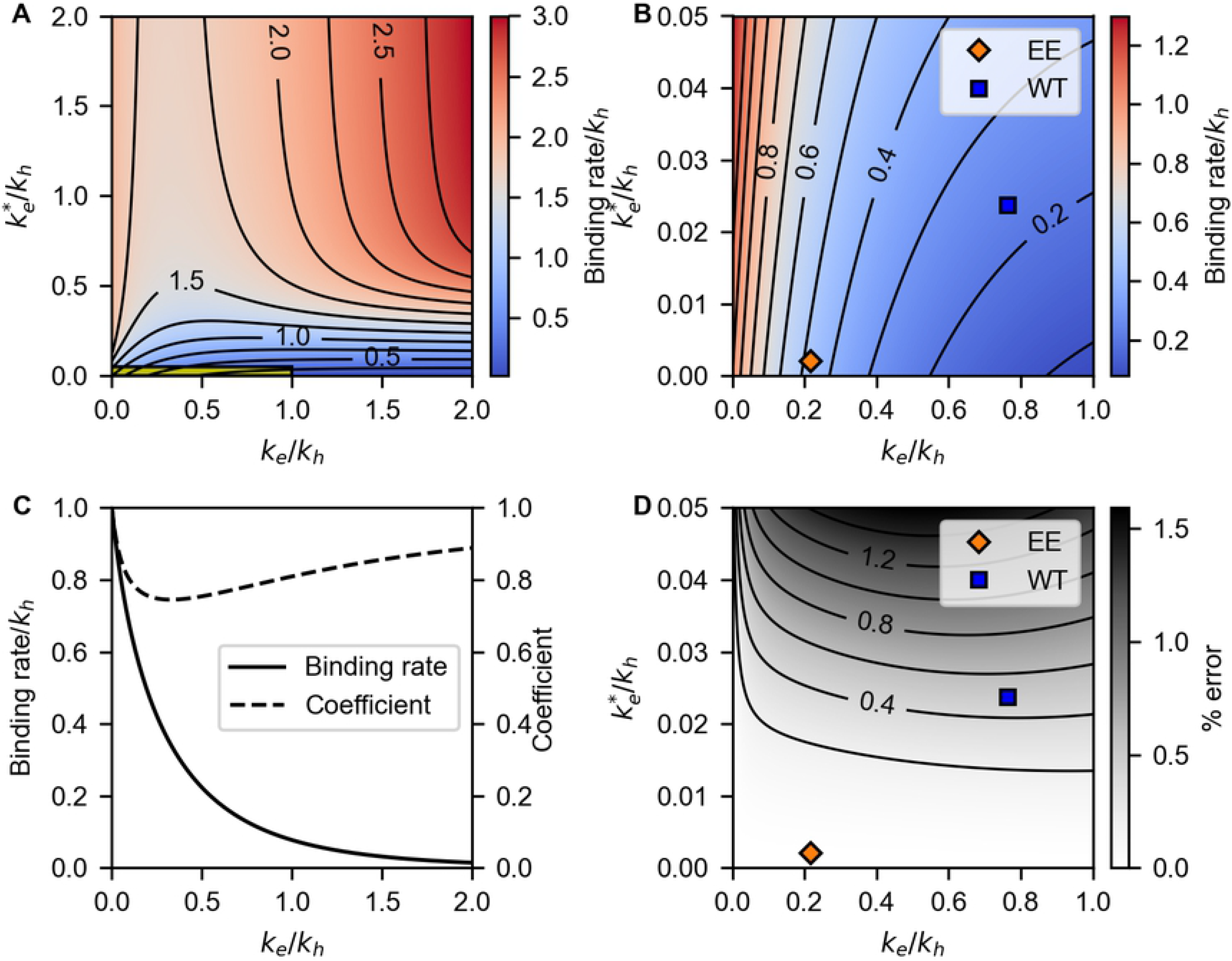
Binding rate of the hexameric model, *γ*_hexa_. All rate constants are normalized by *k*_h_. (A-B) 2D plots of *γ*_hexa_ as a function of *k*_e_ and 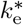. The yellow region in (A) is magnified in (B). Blue and orange squares correspond to the best-fit parameters of KaiC-WT and KaiC-EE. Contours are plotted every 0.25 in (A) and 0.1 in (B). (C) The relaxation rate of Eq (8) *γ*_hexa_ (solid curve) and the coefficient *α* in Eq (9). (D) Percent error of the perturbative approximation of *γ*_hexa_ shown in Eq (10). Blue and orange squares are the same as (B). Contours are plotted every 0.2%.

However, in contrast to the monomeric scheme, *γ*_hexa_ increases with decreasing *k*_e_ when 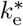 is small. With this property, KaiC-EE of the hexameric model achieves the simultaneous promotion and acceleration of the complex formation by lowering *k*_e_ and 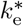 (Fig 2B).

The next problem is why *γ*_hexa_ increases with decreasing *k*_e_ when 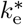 is small. Since 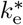 is small, we here approximate *γ*_hexa_ by a perturbation with respect to 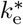 (see Methods). The zeroth-order term of *γ*_hexa_, which we denote by *γ*_hexa,0_, is given by the relaxation rate of

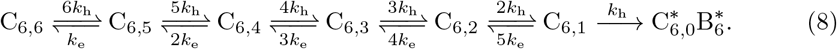

Because the final destination of this scheme is 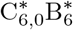, *γ*_hexa,0_ can be regarded as a generalized rate constant from C_6,1−6_ to 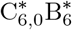. On the other hand, the first-order term of *γ*_hexa_ is given by the relaxation rate of

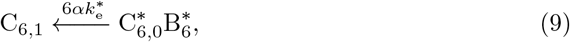

where *α* is the coefficient determined by the perturbation. Thus, *γ*_hexa_ is approximately given by the sum of the generalized rate constants of the forward and backward processes as

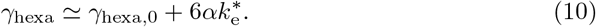

Fig 2D shows that this expression well approximates *γ*_hexa_ with less than 1.5% error around the optimized parameter set. We plot *γ*_hexa,0_ and *α* as a function of *k*_e_ in Fig 2C. Because *α* does not change drastically with *k*_e_, and because 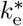 is small, the *k*_e_ dependence of *γ*_hexa_ is mainly determined by *γ*_hexa,0_. Moreover, as shown in Fig 2C, *γ*_hexa,0_ is a decreasing function of *k*_e_ because the increase of *k*_e_ obviously inhibits the transition to 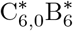 in Eq (8). Therefore, *γ*_hexa_ increases with decreasing *k*_e_ when 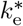 is small.

### Reduction of the ATPase activity of C1 by the complex formation

As a corollary from the KaiB-KaiC complex formation explained above, we here show that the complex formation reduces the ATPase activity, i.e., the ADP production rate, of C1 in the present model. Because the ATPase activity par KaiC monomer *γ*_ATPase_ is represented as

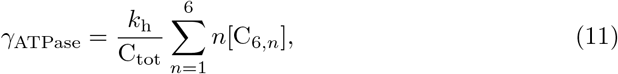

the equilibrium shift in Eq (4) by the complex formation reduces *γ*_ATPase_. Note that the reduction of the ATPase activity has not been explained by any mathematical model so far. The key factor in the present model is Assumption (vi), the difference between the ADP/ATP exchange rate of the binding-incompetent and binding-competent C1. In the present model, the binding of KaiB to C1 stabilizes the binding-competent C1 with a low ADP/ATP exchange rate and thus reduces bound ATP in C1. On the other hand, other previous mathematical models of the KaiABC oscillator do not consider any C1 or KaiB binding dependence of the ADP/ATP exchange rate of C1 [23]. This is why the KaiB binding does not alter the ATPase activity in the previous models.

With the present model and parameters sampled above, we calculate the time courses of the ATPase activity of C1 in the presence of KaiB (Fig 3A). Since the initial state is the steady state of the KaiC only system, the initial ATPase activity corresponds to that in the absence of KaiB. Although the values of the ATPase activity are widely distributed (Figs 3B and C), the reduction by the complex formation is common in all cases (Fig 3A) as explained above.

**Fig 3.**
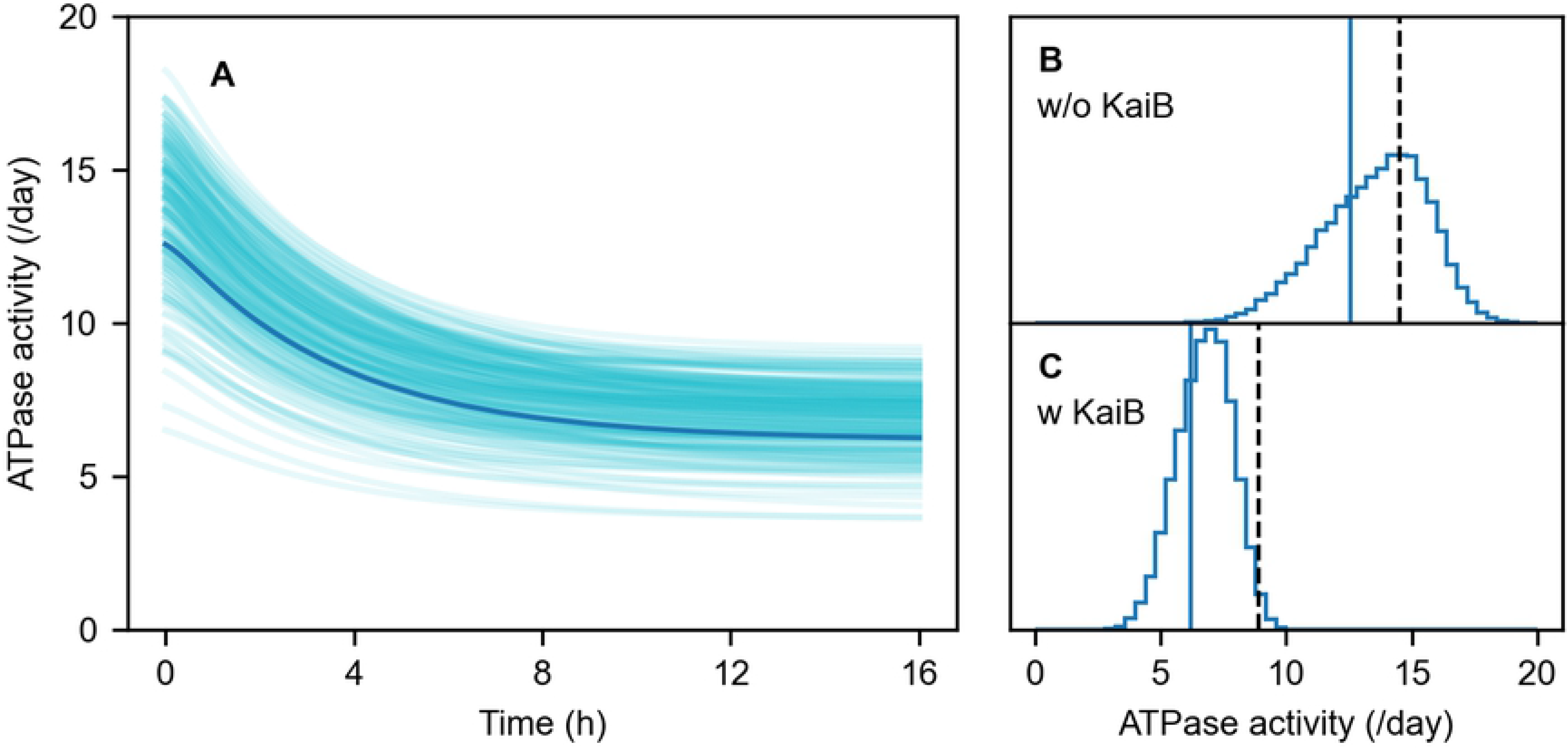
Results of the Bayesian parameter estimation on the ATPase activity of C1. (A) Time-courses of the ATPase activity of C1 during the KaiB-KaiC complex formation. Bold curve corresponds to the best-fit parameters of the hexameric model. Thin solid curves show 200 results of the hexameric model randomly chosen from the MCMC sampling. (B) and (C) Distributions of the ATPase activities in the absence (B) and presence (C) of KaiB. Solid vertical lines correspond to the best-fit parameters of the hexameric model. Dashed vertical lines show the experimental data of the ATPase activities of the whole KaiC.

## Discussion

In the present article, we focused on the simultaneous promotion and acceleration of the KaiB-KaiC complex formation by the phosphorylation of C2, i.e., KaiC-EE has a larger affinity and binding rate of the complex formation than KaiC-WT [16]. This complex formation depends on the nucleotides binding to C1 because the binding-competent conformation of C1 emerges when C1 binds ADP [14, 15, 22]. In particular, this complex formation is considered to be promoted by inhibiting the backward process (ADP/ATP exchange) of the pre-binding processes of C1 rather than by activating the forward process (ATP hydrolysis) [23]. However, inhibition of the backward process generally decelerates the whole process. Thus, without any extension, the previously proposed mechanism cannot describe the acceleration of the KaiB-KaiC complex formation occurring together with its promotion.

To explain the molecular mechanism of the acceleration, we proposed a reaction model that explicitly considers the hexameric form of KaiC in Eq (4). We assumed in this model that six KaiB monomers cooperatively and rapidly binds to C1 with the stabilization of the binding-competent conformation of C1 only when C1 binds six ADP (Assumptions (i-v)). Moreover, to describe the increase of the binding-competent C1 in the presence of KaiB, we assumed that the ADP/ATP exchange rate of the binding-competent C1, 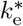, is lower than the binding-incompetent C1, *k*_e_ (Assumption (vi)). Under these assumptions, the small 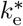 effectively separates the processes in Eq (4) into the principal fast processes from binding-incompetent to binding-competent state (Eq (8)) and the minor slow backward process (Eq (9)). The principal forward processes in Eq (8) consist of the multiple ATP hydrolyses and ADP/ATP exchanges because the binding is assumed to occur only when C1 binds six ADP. In these processes, the ADP/ATP exchange of the binding-incompetent C1 retards the transition to the binding-competent C1. Therefore, the inhibition of this ADP/ATP exchange results in the acceleration of the complex formation.

We also examined a monomeric reaction model in Eq (6), which can be derived from the assumption that the monomers in a KaiC hexamer are independent of each other (Assumption (iv)’). The monomeric scheme consists of the forward and backward processes between the binding-incompetent and binding-competent C1. In contrast to the hexameric model, the forward process is a solo ATP hydrolysis and is irrelevant to the ADP/ATP exchange. Thus, the forward process is never accelerated by lowering the ADP/ATP exchange. This is why the monomeric model cannot describe the simultaneous promotion and acceleration of the complex formation.

The comparison between the hexameric and monomeric models shows that the simultaneous promotion and acceleration of the complex formation arises from the cooperative binding of six KaiB to C1. Our previous study has shown that this cooperative binding originates from the attractive adjacent KaiB-KaiB interaction in the KaiB-KaiC complex [20]. In particular, this KaiB-KaiB interaction much more largely stabilizes the KaiB_6_KaiC_6_ complex than the other smaller complexes KaiB_*n*_KaiC_6_ (1 ≤ *n* ≤ 5), resulting in the predominant formation of the KaiB_6_KaiC_6_ complex. This is because the ring-shaped alignment of KaiB in the KaiB_6_KaiC_6_ complex efficiently acquires the number of adjacent KaiB-KaiB pairs compared to linear alignment. Therefore, we can conclude that the molecular origin of the simultaneous promotion and acceleration of the complex formation is the ring-shaped multimeric structure of KaiC.

It should also be noted that the present reaction model is the first to propose that the ADP/ATP exchange rate of C1 depends not only on the phosphorylation of C2 but on the conformational state of C1 as 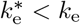. This assumption explains the simultaneous promotion and acceleration of the KaiB-KaiC binding and the reduction of the ATPase activity in the presence of KaiB [24]. These successful results suggest that the conformational state of C1 should be considered separately from that of C2, which highly depends on the phosphorylation of C2, in contrast to other previous models considering only one global conformational change of KaiC [23, 28, 29].

Although the present model explains the simultaneous promotion and acceleration of the complex formation, its significance in the KaiABC oscillator remains elusive. However, for example, it can be expected that this property plays a role in the period robustness against environmental perturbation such as temperature compensation of period. That is, the acceleration of the complex formation due to extensive phosphorylation of C2 may be able to prevent the phosphorylation from taking time by terminating it early. To investigate the functional role, we will extend the present model of C1 to the whole KaiC in future work.

## Methods

### Bayesian parameter estimation

In the framework of Bayesian parameter estimation, the estimation result is represented in the form of the conditional (posterior) probability of the rate constants **k** given the observed experimental data **y**_exp_, which we denote by *p*(**k|y**_exp_). The basic rule of conditional probability known as Bayes’ rule yields an expression of *p*(**k|y**_exp_) by the conditional probability called the likelihood function *p*(**y**_exp_|**k**) and the prior probability of **k**, *p*(**k**):

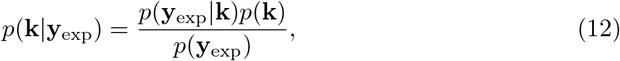

where *p*(**y**_exp_) = *∫d***k***p*(**y**_exp_|**k**)*p*(**k**).

The prior probability *p*(**k**) is determined by a priori knowledge of **k**. Since we only know that *k* is positive and not too large, we assume that *p*(*k*) for each 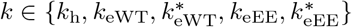 is the uniform distribution on the interval [0, 100] (h^−1^).

We assume for the likelihood function *p*(**y**_exp_|**k**) that, when **k** is given, each experimental data *y*_exp*,i*_ obeys the normal distribution with mean *y*_model*,i*_(**k**) and standard deviation *σ*_*i*_, where *y*_model*,i*_(**k**) is the corresponding value obtained from the model with the given **k**. Thus, the logarithm of *p*(**y**_exp_|**k**) is given by

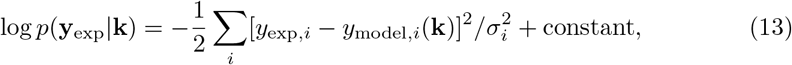

where the index *i* runs over all the time points and the type of KaiC (i.e., KaiC-WT and KaiC-EE) of the data on the KaiB-KaiC complex formation. Although *σ*_*i*_ should also be estimated as well as **k** for rigorous parameter estimation, we fix all *σ*_*i*_ to 0.05 in the present study since the aim is to get a qualitative behavior of the model.

In the parameter estimation of the hexameric model, we additionally consider the ATPase activities of KaiC in the presence and absence of KaiB. Since the experimentally observed values consist of the contributions from both C1 and C2, they can use as upper limits of the ATPase activity of C1 in the present model. Specifically, we add the following term to Eq (13):

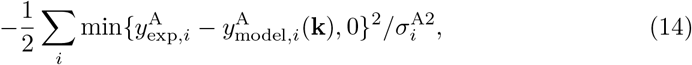

where 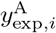 is the experimental data of the ATPase activity of the whole KaiC and 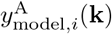 is the model output of the ATPase activity of C1. The index *i* runs over the two cases in the presence and absence of KaiB. For simplicity, we fix 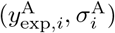 to (8.9, 0.9) (/Day) in the presence of KaiB and to (14.5, 2.0) (/Day) in the absence of KaiB.

We numerically sample the posterior distribution *p*(**k|y**_exp_) defined in Eq (12) by a MCMC method implemented in a Python package emcee (version 3.0.2) [30]. In this method, multiple walkers in **k**-space are required. We first find the maximum point of *p*(**k|y**_exp_) by the optimization routine in SciPy (version 1.5.2). The rate equations are integrated by SciPy with the LSODA method. Then, we prepare 32 walkers with their initial points created by adding random perturbation to the maximum point and sample 50000 points every chain. Following the procedure in the method implemented in emcee [31], we compute the autocorrelation times of the sampled chains *τ*_chain_. Then, we discard the initial 2 max{*τ*_chain_} (= 155) samples from each chain and thin the samples by choosing every 0.5 min{*τ*_chain_} (= 33) sample.

### Perturbative approximation of the binding rate

We here briefly review the perturbative approximation of the eigenvalues of a matrix. For simplicity, we summarize only the case where a matrix *A* is real and diagonalizable with real eigenvalues and eigenvectors. The transition matrix of first-order rate equations belongs to this class. We denote the matrix consisting of the right eigenvectors by *P*, its inverse matrix by *Q*, and the diagonalized matrix of *A* by *D*, that is,

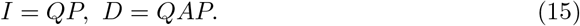

We expand *A*, *P*, *Q*, and *D* with respect to a small parameter *ϵ* like

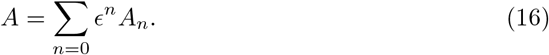

Comparison of the first-order terms of Eq (15) yields

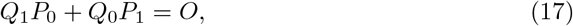

and

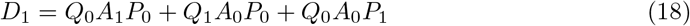

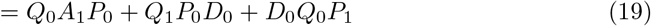

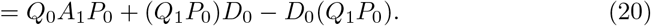

Because the diagonal elements of the second and third terms of the right hand side cancel out each other, the diagonal elements of *D*_1_ coincide with those of *Q*_0_*A*_1_*P*_0_. Thus, the first-order term of the perturbative approximation of an eigenvalue of *A* is given by the “average” of *A*_1_ by the corresponding right and left eigenvectors of *A*_0_.

## Acknowledgements

This work has been supported by JSPS KAKENHI, Grant Number JP18K14185 (SK) and JP21H04676 (SS), and the Indo-Japan bilateral collaboration program. The calculations were partially carried out on computers at the Research Center for Computational Science, Okazaki, Japan.

## Notes

### Competing Interest Statement

The authors have declared no competing interest.

